## Supplementary figures and images for "Boosting Brain Signal Variability Underlies Liberal Shifts in Decision Bias"

### Figure 1 supplement 1

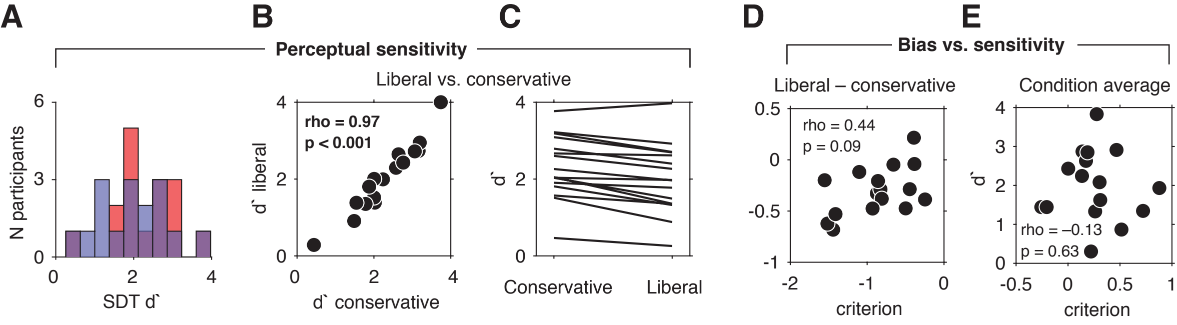

### Figure 3 supplement 1

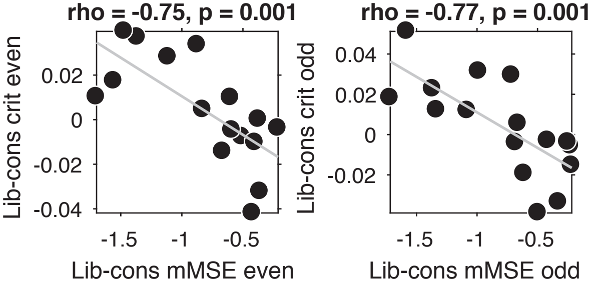

### Figure 3 supplement 2

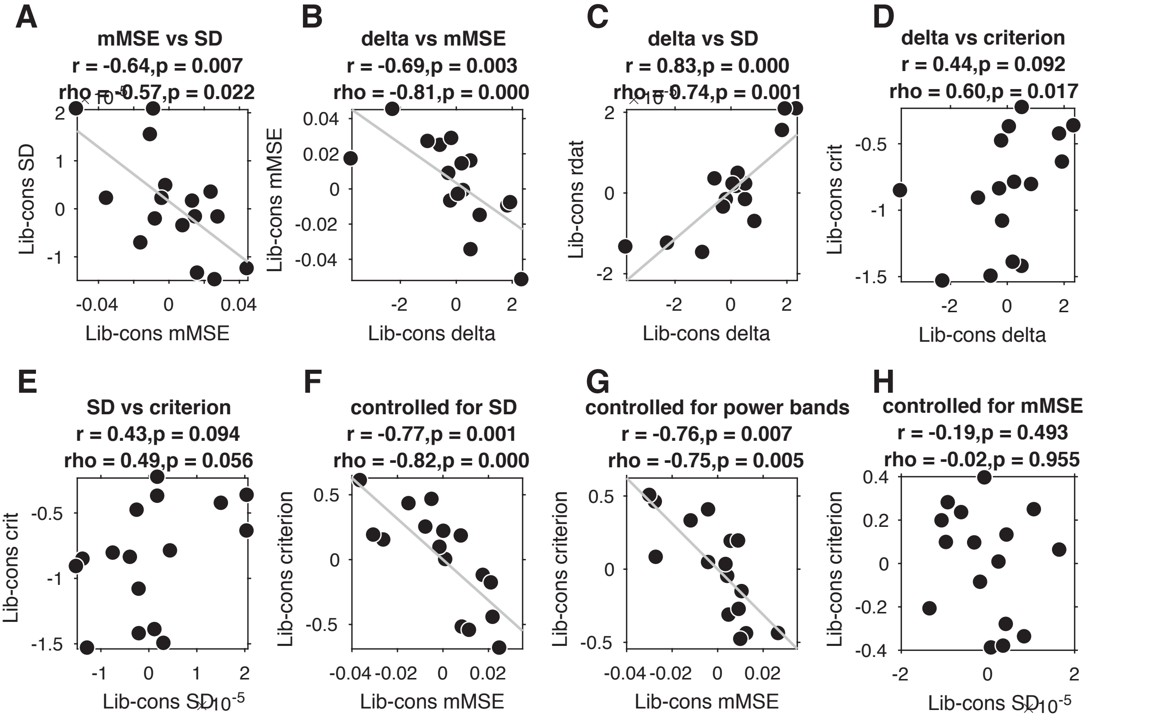

### Figure 3 supplement 3

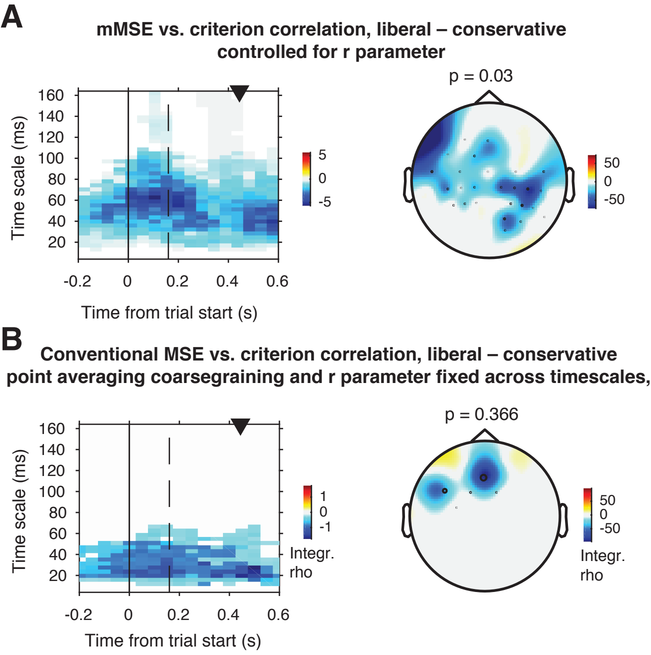

### Figure 3 supplement 4

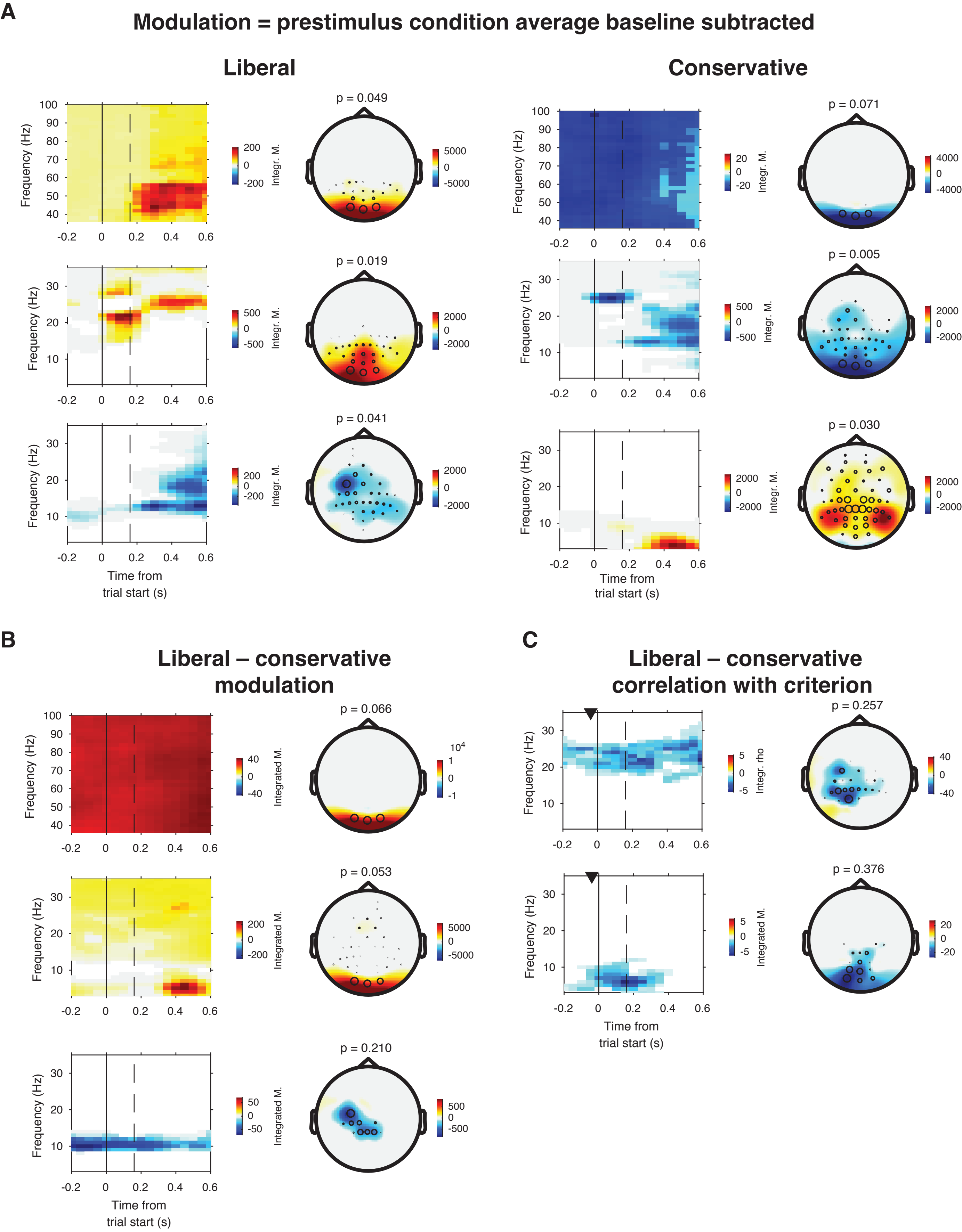

### Figure 3 supplement 5

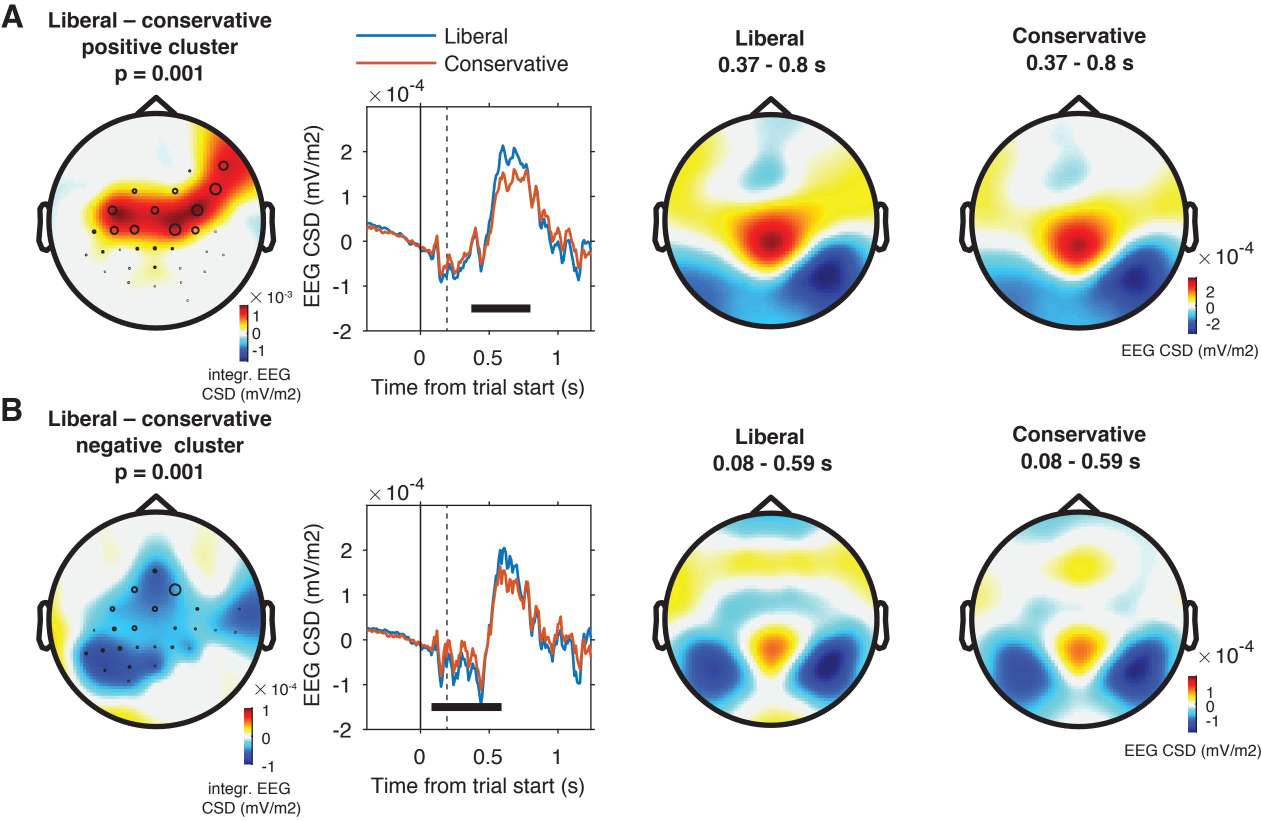
